# The molecular signatures of compatible and incompatible pollination

**DOI:** 10.1101/374843

**Authors:** Chie Kodera, Jérémy Just, Martine Da Rocha, Antoine Larrieu, Lucie Riglet, Jonathan Legrand, Frédérique Rozier, Thierry Gaude, Isabelle Fobis-Loisy

## Abstract

Fertilization in flowering plants depends on the early contact and recognition of pollen grains by the receptive papilla cells of the stigma. To identify the associated molecular pathways, we developed a transcriptomic analysis based on single nucleotide polymorphisms (SNPs) present in two *Arabidopsis thaliana* accessions, one used as female and the other as male. We succeeded in distinguishing 80 % of transcripts according to their parental origins and drew up a catalog of genes whose expression is modified after pollen-stigma interaction. Global analysis of our data reveals that pattern-triggered immunity (PTI)-associated transcripts are upregulated after compatible pollination. From our analysis, we predicted the activation of the Mitogen-activated Protein Kinase 3 on the female side after compatible pollination, which we confirmed through expression and mutant analysis. Our work defines the molecular signatures of compatible and incompatible pollination, highlights the active status of incompatible stigmas, and unravels a new MPK3-dependent cell wall feature associated with stigma-pollen interaction.

## Introduction

In flowering plants, the early interaction between the tip of the female organ (stigma) and the male gametophyte (pollen grain) act as a checkpoint for fertilization. This first step of the female-male interaction includes recognition by the female tissues of the male partner. In the Brassicaceae, sophisticated mechanisms have evolved that allow the papilla cells of the stigma to reject self pollen while accepting non-self pollen. These self/non-self recognition mechanisms are underlie self-incompatibility and promote genetic variability within species. Following compatible pollination, the dry pollen grain starts to hydrate on the stigma papilla and ultimately germinates, producing a tube that penetrates the wall of stigmatic cells and grows down to convey the male gamete towards the ovules for fertilization^1,2^. By contrast, when a pollen grain is recognized as incompatible, it fails to hydrate properly and shows defective germination. This rejection mechanism is initiated by a ligand-receptor interaction and is genetically controlled by a single polymorphic locus called the S-locus^3^. The S-locus Cysteine Rich protein (SCR)/ S-locus protein 11 (SP11) located on the pollen surface interacts with its cognate S-locus Receptor Kinase (SRK) localized to the plasma membrane of the papilla cells^4,5^. This interaction leads to the phosphorylation of SRK that triggers the downstream cascade leading to pollen rejection^1,5,6^. Cellular events triggered in the stigma papillae by these two pathways, compatible and incompatible, have started to be more clearly defined. Compatible pollination induces actin network orientation, calcium export and polarized secretion towards the pollen grain^7–11^. Incompatible pollen leads to inhibition of actin polymerization and vesicular trafficking accompanied by a strong calcium influx within the stigmatic cell^7,12,13^. Stigmatic calcium fluxes were reported to involve the autoinhibited Ca^2+^-ATPase13 (ACA13) for pollen acceptance^8^ and a glutamate receptor-like channel (GLR) for pollen rejection^12^. In addition, the stigmatic EXO70A1 protein was identified as a factor required for polarized secretion during compatible pollination, which is negatively regulated in incompatible reaction^13,14^.

Whereas these results help us understand specific downstream pathways, they are dependent on strong *a priori*. To obtain a global picture of the early fertilization events with no *a priori*, transcriptome approaches were also conducted. The main goal was to draw up catalogues of genes modulated during pollination so as to unravel the stigmatic response to compatible or incompatible pollen^8,15–18^. However, the main drawback of these approaches was the difficulty in accurately distinguishing between pollen and stigma derived transcripts. To address that issue, translatome analysis^19^ has recently been applied to identify sex-specific genes expressed during pollination, but this strategy needs great amounts of tissues from the specific transgenic line and fine techniques of biochemistry.

Here, inspired by previous RNA-seq analysis^20^, we developed a new experimental procedure coupled with a bioinformatic analysis of sequencing data to comprehensively unravel the dynamic events that occur both in the stigma and pollen grain following compatible and incompatible pollinations. We took advantage of the SNPs existing between two distinct *Arabidopsis thaliana* accessions, one used as female (Col-0) and the other as male (C24), to differentiate male and female transcripts based on a new statistical methodology, ASE-TIGAR^21^ which can take all sequenced reads in account even those without SNPs while previous study used only reads with SNPs^20^. Our analysis allowed the identification of 80 % of mRNAs according to their parental origin and revealed transcriptional changes occurring specifically in either the stigma or pollen grain/tube. Gene Ontology and signaling pathway prediction of up-regulated genes showed that pattern-triggered immunity (PTI) pathways, including induction of the Mitogen-activated Protein Kinase 3 (MPK3), were activated on the female side after compatible pollination. Mutant analysis then confirmed that MPK3 is implicated in the growth of pollen tubes in papilla cells.

## Results

### Experimental setup to isolate transcripts from compatible and incompatible pollination in *A. thaliana*

To compare self-compatible and self-incompatible pollination, we needed to have the same genetic background, albeit with different stigmatic responses to pollen. *Arabidopsis thaliana* has a high level of self-fertilization due to mutations that disrupt the self-incompatibility system present in its outbreeding ancestor^22^. To obtain a self-incompatible *Arabidopsis thaliana* background, we transformed *A. thaliana* with a functional *SRK-SCR* gene pair isolated from its close self-incompatible relative *A. lyrata*^23,24^. To analyze both compatible and incompatible reactions and take advantage of nucleotide polymorphisms between *A. thaliana* accessions, we generated two transgenic lines that restored the incompatible response: one in the Col-0 background expressing the *SRK* gene from the *A. lyrata S*14 haplotype (Col-0/*SRK14*) and the other in C24 background expressing the *SCR* gene from the same *S*-haplotype (C24/*SCR14*) (Fig. 1a, Supplementary Fig. 1a). To express the SRK protein in stigmatic cells, we used the *SLR1* promoter that displays a strong stigma-specific activity in *Brassicaceae*^25,26^. Expression of the *SCR* gene was controlled by its own promoter. We selected one transgenic line for Col-0/*SRK14* and one for C24/*SCR14*. Both lines were self-fertile.

**Fig. 1.**
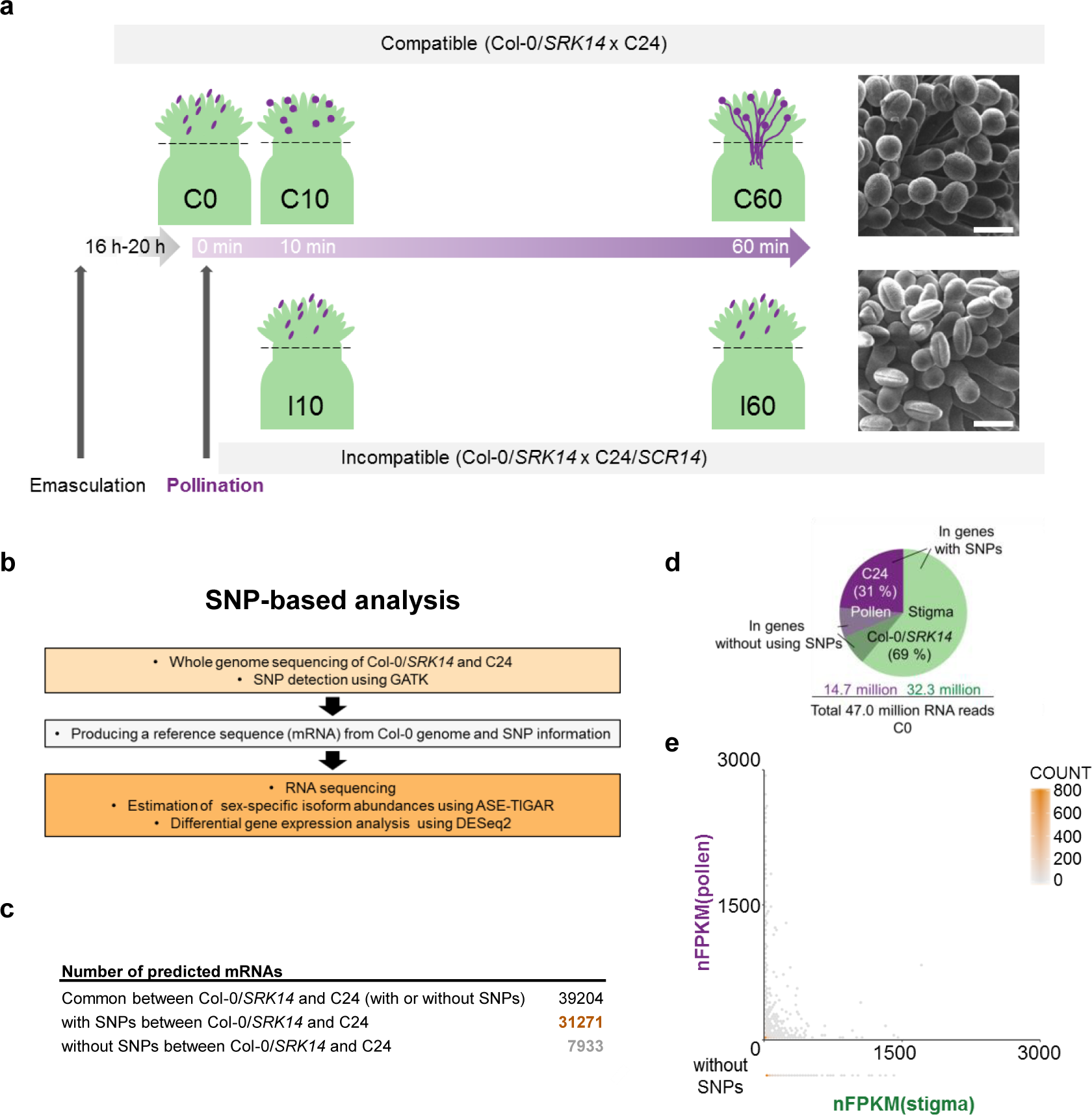
SNP-based RNA-seq analysis. **a**, Time course of sample collection. Flowers of Col-0/*SRK14* were emasculated 16h - 20h before pollination with compatible (C24) or incompatible (C24/*SCR14*) pollen grains, respectively. Stigmas were harvested just before the stigma (dashed line) 0, 10, 60 min after compatible (C0, C10, C60) or incompatible (I10, I60) pollination for RNA extraction. Typical scanning electron microscope images for compatible (Col-0 /*SRK14* x C24) and incompatible (Col-0 /*SRK14* x C24/*SCR14*) reaction observed 60 min after pollination. Scale bar = 20 μm. **b**, Three steps of a SNP-based data analysis pipeline. **c**, Number of predicted mRNAs derived from the reference sequence produced at the second step in b. Around 80 % of coding regions have SNPs between Col-0/SRK14 and C24. **d**, Number and ratio of estimated read counts for Col-0/*SRK14* and C24 at C0 computed by ASE-TIGAR at the third step in b. Reads estimated to the stigma or pollen include reads assigned to the genes without using SNPs. **e**, Hexibin plot of nFPKM for Col-0/*SRK14* and C24 at C0. nFPKM(stigma) of genes without SNPs were plotted outside.

Next, we tested the fertilization response in crosses between these lines. As expected, stigmas of Col-0/*SRK14* pollinated with wild-type C24 pollen showed a clear compatible reaction with hydration of pollen grains and more than 55 pollen tubes elongation on papilla cells (Fig. 1a upper part, Supplementary Fig. 1bd). By contrast, when Col-0/*SRK14* stigmas were pollinated with C24/*SCR14* pollen, a strong incompatible reaction was observed as deduced from poor pollen hydration and the absence of pollen tube germination (Fig. 1a lower part, Supplementary Fig. 1cd).

Building on these experimental validations, we performed a time-course experiment focusing on two time-points of the interaction to identify genes whose expression is modified following pollen-stigma interaction. We selected an early stage (10 minutes after pollen deposition), which corresponds to the start of pollen grain hydration, and a later stage (one hour after pollination) when pollen tubes reach the base of the stigma. We sequenced mRNAs extracted from pollinated stigmas at 10, and 60 min after compatible (Col-0/*SRK14* x C24) pollination (C10, C60, respectively) and for incompatible (Col-0/*SRK14* x C24/*SCR14*) pollination (I10, I60, respectively) (Fig. 1a). We also extracted mRNA right after pollination, 0 min (Col-0/*SRK14* x C24), as a control for both compatible and incompatible pollination (C0).

### SNP-based transcriptome analysis using variants between Col-0 and C24

To distinguish the parental origin of the transcripts, we developed a method based on the detection of small genomic variations, including SNPs and small insertions and deletions (indels) between Col-0 and C24 accessions. The pipeline includes three main steps (Fig. 1b, Supplementary Fig. 2). In the first step, variations between Col-0/*SRK14* and C24 genomes were identified by whole-genome sequencing of the two strains with the read depth of 9.3 X for Col-0/*SRK14* and 14.2 X for C24 (Supplementary Table 1). After aligning clean reads against the reference genome (TAIR10, accession Col-0), SNPs and short indels between the mapped reads and the reference genome sequence were identified. Reads from Col0/*SRK14* and C24 covered 95.8% and 90.6% of TAIR10 genome sequence, respectively. When compared to the reference, the line Col-0/*SRK1*4 had 2,032 variants, while C24/*SCR14* had 732,767 variants, respectively. In a second step, we introduced previously identified variants in the sequence of the TAIR10 template to produce two new reference genome sequences, one for Col-0/*SRK14* and one for C24 (Fig. 1b, Supplementary Fig.2). At the end of this process, between the new genomes of Col-0/*SRK14* and C24, we identified 616,781 SNPs and 446,999 indels (Supplementary Table 2). 24 % and 31 % of genomic polymorphism were found in UTRs and CDSs for Col-0/*SRK14* and C24, respectively, and led to sequence variations in predicted mRNAs (Supplementary Table 2). We then annotated these two new genome sequences by projecting gene models from TAIR10. We predicted 39,205 gene models in Col-0/*SRK14* and 39,206 in C24. We then extracted predicted mRNA sequences for each gene model from these annotated Col-0/*SRK14* and C24 genome sequences to produce maternal and paternal reference transcripts. We obtained a list of 39,204 predicted gene models that were shared between maternal and paternal reference. The number of predicted mRNAs with at least one SNP between Col-0/*SRK14* and C24 was 31,271 among the total common predicted mRNAs (39,204) (Fig. 1c). This result allowed us to distinguish the origin of about 80 % (31,271/39,204 = 79.8%) of mRNAs with SNP-based analysis. The last step included RNA read mapping and the estimation of sex-specific isoform abundances using ASE-TIGAR. The total length of raw reads from each condition was more than 6,400 Mb (Supplementary Table 3). ASE-TIGAR uses a Bayesian approach to estimate allele-specific expression^21^. This approach allowed us to use all sequenced reads even those without SNPs. After obtaining the estimated read counts from ASE-TIGAR, we used DESeq2^27^ to normalize counts, for female and male transcripts, respectively (Fig. 1b, Supplementary Fig. 2 Supplementary Table. 4).

**Fig. 2.**
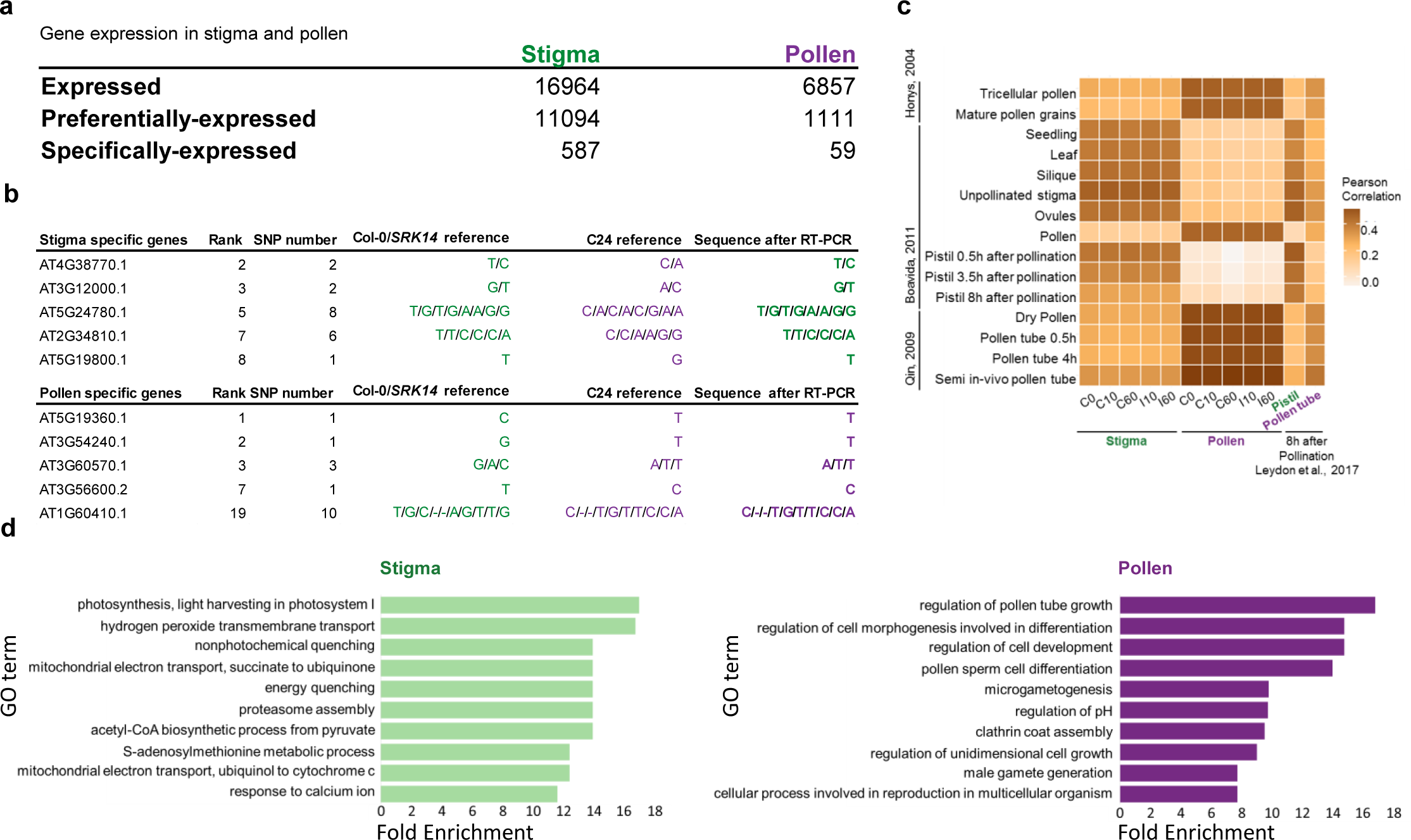
Validation of the SNP-based analysis. **a**, Number of stigma or pollen (preferentially / specifically) -expressed genes at C0 selected by nFPKM (Supplementary Table 5). **b**, RT-PCR and sequence analysis of stigma or pollen specifically-expressed genes at C0. The genes are selected among the 20 specifically-expressed genes and the rank of the genes according to the expression level are presented. RNA was extracted from pollinated stigmas at C0 then we analysed their SNP-information by RT-PCR and sequencing. SNP number corresponds to the number of SNPs present in the sequenced regions. **c**, Heat map of Pearson’s correlation coefficient between the stigma/pollen transcripts from the SNP-based analysis and the previously published transcripts. **d**, Top ten gene enrichment categories of stigma or pollen preferentially-expressed genes at C0 according to the GO term on biological processes. Enrichment analysis was performed with the one thousand top expressed genes in stigma (left) or pollen (right) among the list, respectively. Only top 10 first enrichment categories are shown. Selection criteria for genes analysed in **b** (sex-specific), and **d** (sex-preferential), are described in the manuscript.

From this large dataset, we first determined the contribution of each tissue (stigma vs pollen) in mixed samples harvested immediately after compatible pollination (C0). We found that among the 47.0 million RNA reads, 69 % were estimated as derived from Col-0/*SRK14* (stigma), and 31 % were from C24 (pollen) (Fig. 1d) although 15 % of reads were assigned to the genes without using SNPs. This proportion is stable over time (0, 10 min, 60 min) and independent of the pollination type (compatible, incompatible), thus suggesting that these proportions are reflecting the relative abundance of stigmatic cells compared with pollen grains collected by our manipulation.

Then, we analyzed the relative abundance of each transcript between stigma and pollen. To do so, transcript abundance was computed by ASE-TIGAR and was expressed as Fragments Per Kilo base of exon per Million reads mapped (FPKM), which accounts for sequencing depth and gene length. We then normalized FPKM (nFPKM) by dividing the FPKM of each transcript by the ratio of the transcript counts from stigma [nFPKM(stigma)] or pollen [nFPKM(pollen)] at C0 (Fig. 1d, Supplementary Table 4). Values of nFPKM were displayed as a hexbin^28^ plot to visualize the distribution of gene expression levels. Genes without SNPs did not show any particular pattern (Fig. 1e, bottom line). Gene expression levels were widely distributed and genes highly expressed in female or in male were clearly separated, suggesting the existence of distinct transcript signatures between stigma and pollen (Fig. 1e).

### Post-validation of the SNP-based analysis

To have a global view of expressed genes in stigma and pollen, we constructed three classes of genes from calculated nFPKM: expressed genes, sex-preferentially and sex-specifically expressed genes (see methods for precise criteria of gene selection) (Supplementary Table 5). Briefly, we defined genes that showed nFPKM >1 as expressed genes, those that were expressed at least a ten-fold higher in stigma than in pollen as stigma-preferentially expressed genes, and those that were at least one hundred-fold higher expressed in stigma than in pollen as stigma-specific expressed genes (and vice versa for pollen preferentially and specifically expressed genes) (Fig. 2a).

To evaluate the accuracy of our SNP-based analysis, we first focused on sex-specific genes and analysed the top 20 highly expressed genes (Supplementary Table 5) using the ThaleMine database (https://apps.araport.org/thalemine/begin.do). Heat maps of gene expression levels based on Cheng et al., 2016 were generated (Supplementary Fig. 3). Interestingly, stigma specific and highly expressed genes were clearly excluded from pollen even though many of them were also expressed in various tissues. By contrast, most pollen specific and highly expressed genes showed expression restricted to pollen and stage 12-inflorescences, which contain mature pollens (Supplementary Fig.3) Moreover, within the list of stigma-specific genes, we found *AtS1* (*AT3G12000.1*), which has been described as specifically expressed in *Arabidopsis* stigmas^29,30^, and in the list of pollen-specific genes, we found *CPK34* (*AT5G19360.*1), the protein product of which is involved in pollen tube regulation^31^. Finally, to confirm the specific expression of these genes in stigma or pollen, we carried out RT-PCR and sequence the cDNAs of SNP-containing regions using C0 sample as template. From the top 20 highly expressed genes, we selected genes whose location of SNPs permitted designing of primers. We found that, as expected, mRNAs of genes in the stigma-specific genes had SNPs only from Col-0/*SRK14* (stigma), whereas mRNAs of genes in the pollen-specific genes had SNPs only from C24 (pollen) SNPs (Fig. 2b).

**Fig. 3.**
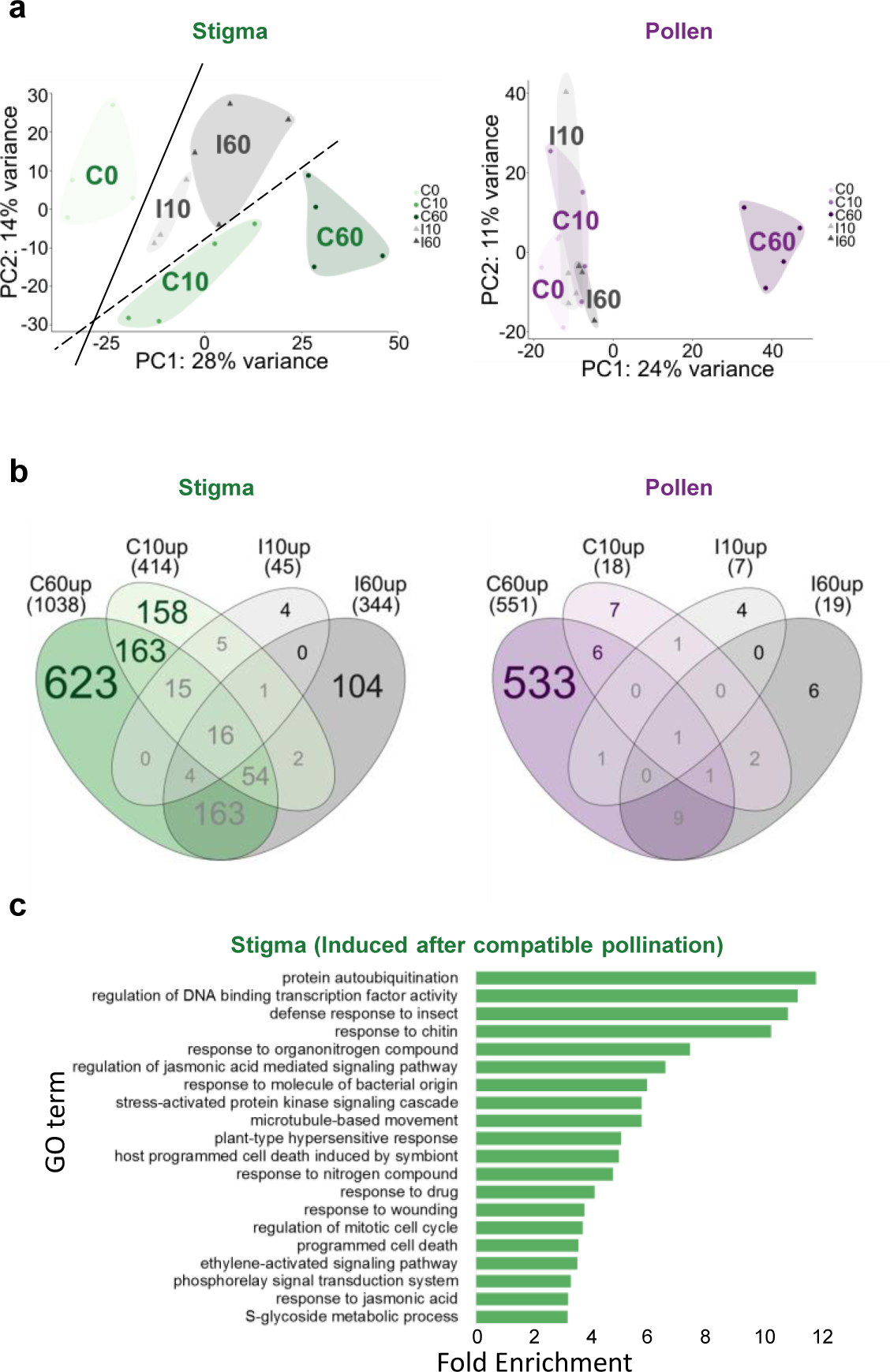
Gene expression dynamics after pollination. **a**, PCA of stigma (left) and pollen (right) transcripts. Biological replicates of all the conditions were analysed. **b**, Venn diagrams showing the number of up-regulated (FC > 2) genes in stigma (left) and pollen (right), after compatible (green and violet) or incompatible (grey) pollinations. **c**, Gene enrichment categories of up-regulated genes after compatible pollination in stigma according to GO term on biological processes. Only genes specifically induced after compatible reaction (b, left; 158+163+623) were analysed. Only top 20 enrichment categories are shown.

To assess the global consistency of our data sets with already published studies, we further analyzed our SNP-based data by correlating them tissue-specific transcripts reported from microarray experiments^32–34^. Stigma associated transcripts from our analysis showed poor correlation with male transcriptomes from mature pollen or growing pollen tube, whereas the highest correlation was observed with transcriptome from unpollinated stigmas. Conversely, our pollen associated transcriptome showed a high correlation with male transcriptomes and only poorly correlated with female transcriptomes or transcriptomes from vegetative tissues (Fig. 2c). Correlations between another recent SNP-based analysis, which identified pistil- and pollen tube-specific transcripts 8 hour after pollination^20^ and the tissue/organ-specific transcripts showed similar trends to our stigma- and pollen-transcripts (Fig. 2c).

To further characterize the pollen vs. stigma associated transcripts, we looked for Gene Ontology (GO) term enrichment at C0 within the list of sex-preferentially expressed genes (FDR < 0.05). Although GO terms may be somewhat subjective or not fully consolidated by functional tests, with the current annotation, we observed a clear difference between the two sets of transcripts (Fig. 2d). From GO term enrichment of the top 1000 highly expressed genes among stigma or pollen -preferentially expressed genes (Supplementary Table 5, Supplementary Table 6), our analysis revealed high enrichment of several GO terms associated with metabolism in stigmas (such as “photosynthesis”, “mitochondrial-”, and “-metabolic process”) suggesting an active metabolic state of stigmatic cells (Fig. 2d left). Conversely, GO terms on the pollen side were specific to pollen functions such as “pollen tube growth”, “pollen sperm cell differentiation” and “cell tip growth” (Fig. 2d right).

Osaka et al.^35^ previously reported the transcriptome of unpollinated stigmas using laser microdissection of *Arabidopsis* stigmatic cells. Among the top 100 expressed genes in this latter analysis, 44 were common with the top 100 expressed genes in stigma of our analysis at C0^35^.

Altogether, these results validate our SNP-based workflow, which allows identification of female- and male-derived transcripts from a combination of tissues following pollination.

### Gene expression dynamics triggered after pollination

To examine the transcriptomic response of pollen and stigma following compatible or incompatible pollination, we first performed a principal component analysis (PCA) ^36^ using the gene expression levels of each biological replicate in all conditions (Fig. 3a). Data from stigma- and pollen-transcripts were treated separately (Fig. 3a left and right, respectively). The total explained variance of the first two principal components (PC1 and PC2) is around 39% (28 +11) for stigma and 35% (24 + 11) for pollen. These relatively low percentages are explained by the fact we took all genes and not only those that are specific for compatible or incompatible reaction. It also explain the fact that the different conditions (C0, C10, C60, I10 & I60) were not always well separated along those axis.

Nevertheless, comparing the PCA of stigma and of pollen transcripts suggested different dynamics in each tissue. On the stigma side, the PCA showed that both compatible and incompatible pollinations triggered a transcriptional response. Along the PC1 axis, capturing almost 30% of the explained variance, samples are temporally sorted from 0 min to 60 minutes, thus suggesting that PC1 can be interpreted as a time-axis. For the second axis, explaining 14% of the observed variability, compatible samples are located in the lower part when incompatible samples are located in the upper part, close to those of the starting point (C0). This seems to indicate that this axis is oriented by the compatible/incompatible effect. Meanwhile, on the pollen side, all the conditions except C60 were clustered together. Again PC1 captures the temporality of the compatible response, however we cannot conclude that PC2 is related to the compatible response effect since early compatible and incompatible clusters overlap.

The clear response pattern of pollen at 1 hour is coherent with the massive changes displayed by compatible pollen, which hydrated and germinated a pollen tube growing in the stigmatic tissue (Fig. 3a).

After performing differentially expressed gene (DEG) analysis, we found more up regulated genes than down regulated genes, and more DEGs in stigma than in pollen (1841 up in stigma vs. 595 in pollen, 513 down in stigma vs. 113 down in pollen) (Supplementary Table 7). To get a clearer picture on the number of genes whose expressions were modified during the course of pollination, we then used a Venn diagram representation (Fig. 3b) only with up regulated genes (padj < 0.1, FC > 2) to study. The Venn diagram of up-regulated genes in stigma (Fig. 3b left) showed similar dynamics to those observed in PCA, with a moderate alteration of gene expression in incompatible reactions, with only 45 genes at I10 and 344 genes at I60. By contrast, following compatible pollination, a massive and progressive change of gene expression was detected with 414 up-regulated genes at C10 and 1038 at C60 (Fig. 3b left). On the pollen side (Fig. 3b right), very few genes showed altered expression, except in condition C60, where 551 genes were up-regulated (6 common in C10 and C60, 533 exclusively in C60) (Supplementary Table 8). Again this correlates perfectly with the PCA. Although the pollen is completely changing its physiology during compatible pollination, the stigma does not remain passive and undergoes a massive molecular reprogramming.

### PTI pathways induced after compatible pollination in stigma

The rapid and global transcriptomic changes in stigma after compatible pollination motivated us to investigate the molecular events occurring at this stage. The 944 genes (158 + 163 + 623) induced only after compatible reaction (Fig. 3b left) showed dramatically different GO terms and enrichment from those preferentially expressed at C0 in the stigma (Fig. 2d left, Fig. 3c). There were 126 terms with high confidence (FDR < 0.05) and 85 terms with FDR < 0.005 (Supplementary Table 9). In particular, we found many GO terms including “response” and “signalling”, thus suggesting the prompt activation of signalling pathways in response to the pollination event. We also applied the enrichment analysis for 108 genes (104 + 4) that were induced only after incompatible pollination in stigma. But this was not informative when compared with compatible reaction as only 9 terms showed a FDR < 0.05 with the minimum FDR = 0.006.

To predict induced pathways after pollination, we then mapped the 944 up-regulated genes after compatible pollination to KEGG pathways^37^. We found many metabolism related pathways, signaling pathways including hormone-signaling (with 20 genes), plant-pathogen interaction (with 22 genes) and MAPK pathways (with 11 genes). Interestingly, two plant-pathogen interaction pathways known as pattern-triggered immunity (PTI) induced by bacterial flg22 and EF-Tu^38^ were clearly distinguishable with 5 up-regulated genes in compatible situation (Fig. 4, green). On the other hand, among the 108 (4+104) exclusively up-regulated genes after incompatible pollination, we could not distinguish clear induction of PTI pathways as only one gene, *EFR*, was up-regulated (Fig. 4, gray).

**Fig. 4.**
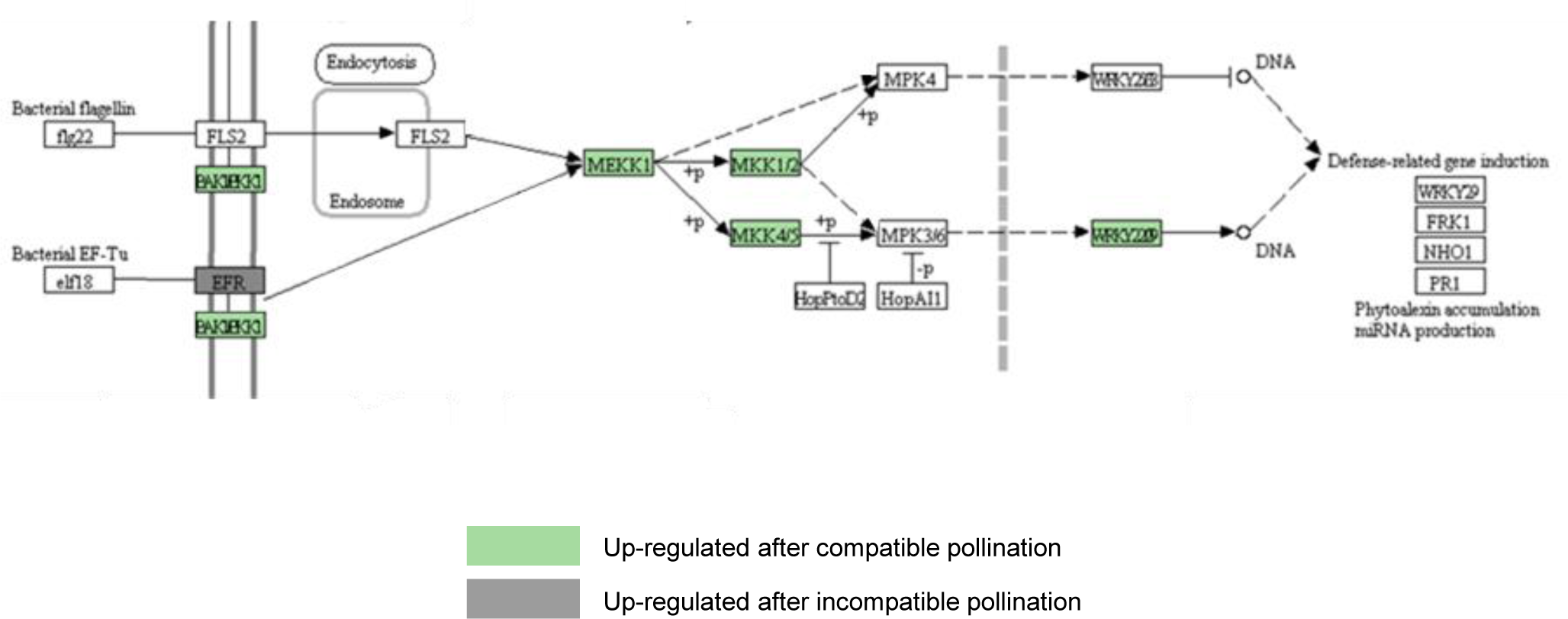
Induced genes in PTI pathways after pollination in stigma. KEGG pathway mapping applied for the genes exclusively up-regulated after compatible pollination (Fig.3 b, left; 158+163+623) and incompatible pollination (Fig.3 b, left; 4+0+104), respectively. Green colored genes were identified from the compatible gene set and grey were from the incompatible set. Up-regulated gene in our analysis assigned to *BAK1*/*BKK1* was *SERK4*, and to *MKK1/2* was *MKK6* (see Fig. 5 and Supplementary Table 10).

In the compatible pollination, *SERK4*, *BAK1*, *EFR* and *FLS2* - all genes known to encode receptors of bacterial components - were expressed in stigma at C0 and *SERK4* was up regulated at C60 (FC = 2.3). Although *MKK6* and the transcription factor-encoding genes *WRKY22* and *WRKY25* were not expressed at C0, the first two were already induced at C10 and induction was maintained at C60, while *WRKY25* was induced only at C60 (all FCs were between 2 and 2.8). *MPK4* and *MPK6* were highly expressed from C0 to C60 compared with other components of the pathway but did not exhibit significant expression change. Interestingly, even though *MPK3*, *WRKY33* and *WRKY53* did not have SNPs, their high induction following compatible (FC = 6.3, 10.0, 11.0 respectively) compared with incompatible (FC = 1.5, 2.8, 5.2) reaction suggests that this PTI pathway is a hallmark of compatible pollination (Fig. 5, Supplementary Table. 10). Altogether, these analyses indicate that the PTI pathway is rapidly induced in stigma only after compatible pollination.

**Fig. 5.**
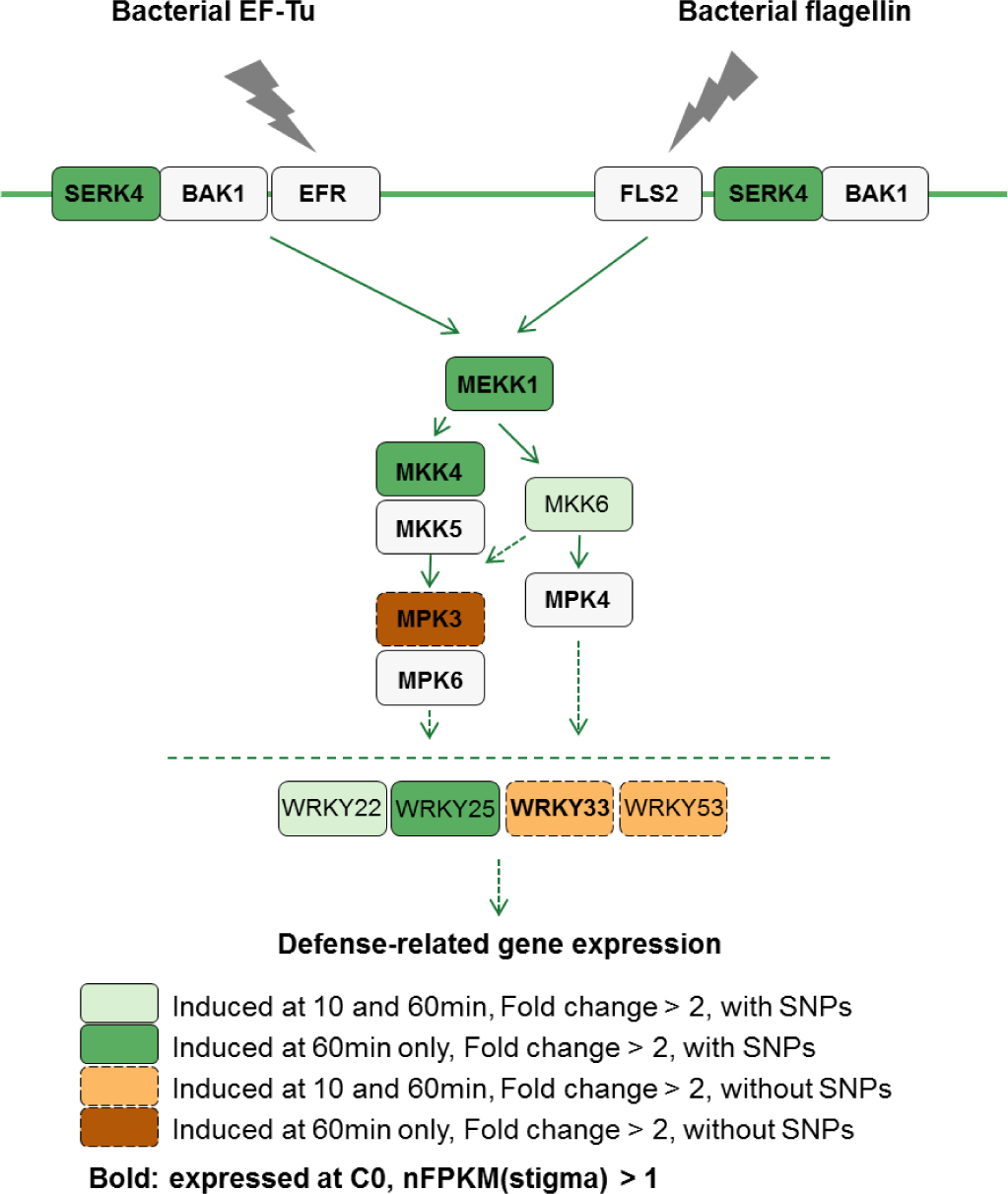
Induced pathways in stigma after compatible pollination. Genes in green boxes with solid line were with SNPs and up-regulated (FC > 2) at 10 min and 60 min (light green), or only at 60 min (green), genes in orange boxes with dashed line were without SNPs and up-regulated (FC > 2) at 10 min and 60 min (orange), or only at 60 min (dark orange). Genes written in bold were expressed at C0; nFPKM(stigma) > 1. The pathway map from KEGG database was modified with the information from Bigeard et al. 2015.

### Stigma-pollen interaction activates and requires MPK3

The predicted mRNA of *MPK3* did not have SNPs between Col-0/*SRK14* and C24, preventing the accurate assignment of the increased expression to stigma tissues. To check whether activation of MPK3-associated pathways occurs in stigma, as predicted from our analysis, we derived the list of genes known to be co-expressed with MPK3 from ATTED-II (http://atted.jp/)^39^. Genes in the list were ordered based on Pearson’s correlation coefficient with MPK3 expression (from high to low) (Supplementary Table 11). After removing the genes without SNPs, we checked the expression of the top 50 genes using our SNP-based analysis in stigma (Supplementary Fig. 4, upper plot) and pollen (Supplementary Fig. 4, lower plot). At C0, most of the genes were moderately expressed in stigma but not in pollen. After compatible pollination, they were induced only in stigma, supporting our hypothesis of a stigma-specific MPK3 pathway induction. Subsequently, we took advantage of the MPK3-knockout line *mpk3-DG*^40–42^ to monitor the expression and activation of MPK3 protein *in vivo*.

Immunoblotting was performed on protein extracts of pollinated stigmas from the following combinations of crosses: Col-0 x *mpk3* (i.e., MPK3 protein only present in stigma), *mpk3* x Col-0 (i.e., MPK3 only in pollen), Col-0 x Col-0 (MPK3 in both), and *mpk3* x *mpk3* as a negative control. Pollinated stigmas were harvested immediately or 60 min after pollination, corresponding to C0 and C60, respectively. The absence of a specific band for MPK3 from the *mpk3* x *mpk3* sample confirmed the complete knockout of the mutant (Fig. 6a). MPK3 protein was detected in both stigma (Col-0 x *mpk3*) and pollen (*mpk3* x Col-0) but with a higher abundance in the stigma (Fig. 6ab). MPK3 was slightly accumulated 60 min after pollination in stigma and in pollen (FC = 1.6 and 2.4, respectively). Subsequently, we checked whether MPK3 was activated through phosphorylation using an anti-Phospho MPK antibody. At C0, the band corresponding to phosphorylated MPK3 was very weak, but its intensity clearly increased after compatible pollination in stigma but not in pollen (FC = 5.2 and 0.9 respectively) (Fig. 6cd). Altogether these results demonstrate that MPK3 is both transcriptionally and post-translationally activated in the stigma during compatible pollination.

**Fig.6.**
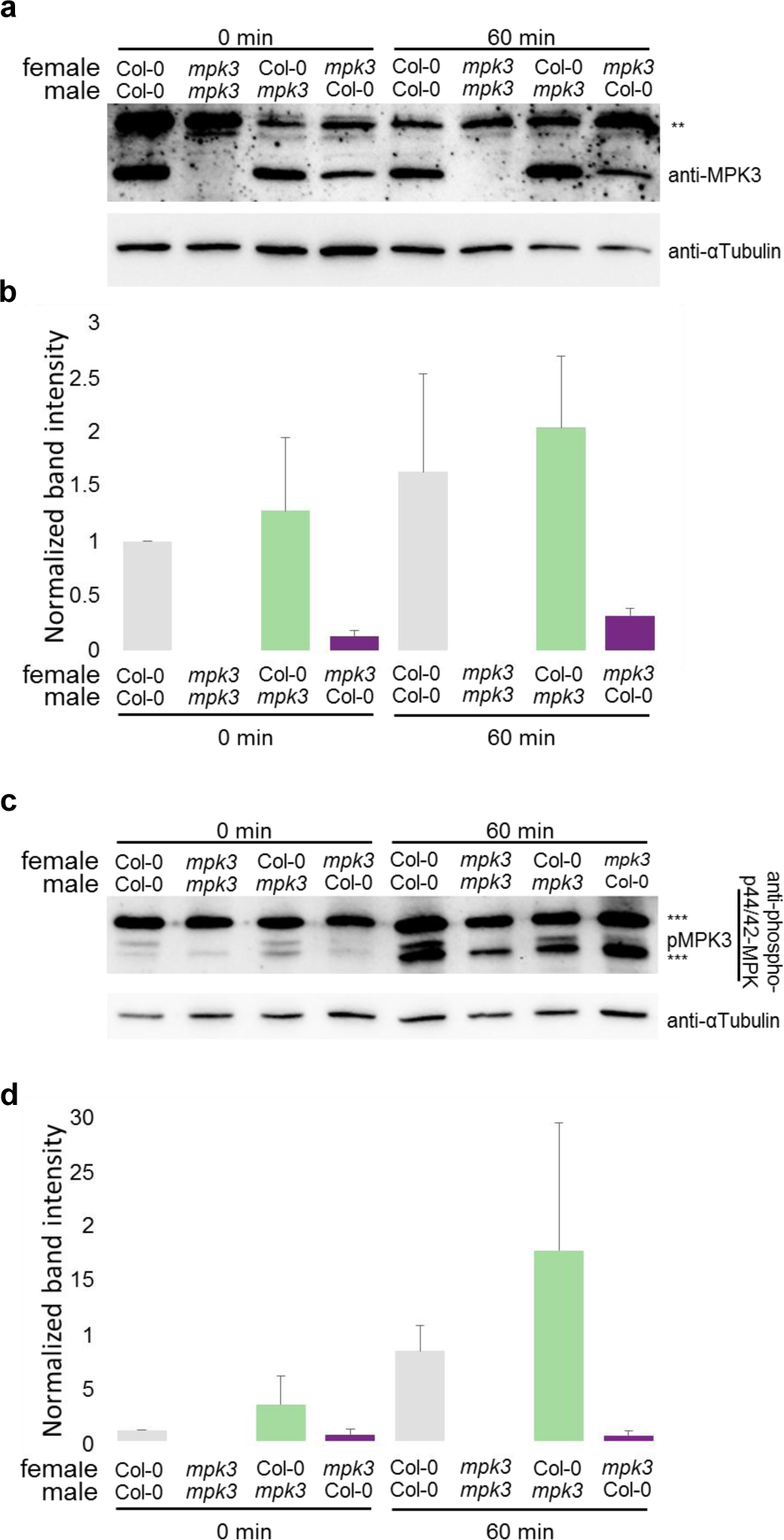
Activation of a MPK3-dependent pathway after compatible pollination. Immunoblotting assays after compatible pollination with *mpk3* mutant line. Pollination between Col-0 x Col-0, *mpk3* x *mpk3*, Col-0 x *mpk3*, and *mpk3* x Col-0 were performed. Protein was extracted from pollinated stigmas at C0 and C60 then analysed. Anti-MPK3 antibody was used for **a**, and anti-phospho-p44/42 MPK antibody was used for **c**. The shown western blots are representative of three independent biological replicates. All of them showed similar results. **b**., **d**. Semi-quantification of the band intensity (b for a, d for c). Green, violet and grey bars indicate signals from stigma side, from pollen side, and from both sides, respectively. Error bars indicate standard error of the mean, ** indicates non-specific signal, *** indicates non-specific signal or signals from other MPK proteins.

To elucidate whether the loss of MPK3 function might have any effect on pollen-stigma interaction, we pollinated *mpk3* stigmas with Col-0 pollen and observed the pollinated stigmas 60 min after pollination by scanning electron microscopy. Col-0 x Col-0 pollination was used as a control. Before pollination, there were no apparent differences between stigmas of *mpk3* and Col-0 (Supplementary Fig. 5). Hydration of pollen grains and elongation of pollen tubes occurred normally on *mpk3* stigmas compared to the control. However, when we measured the width of the contact sites between the pollen tube and stigma (white arrows on Fig. 7a), even if the distributions of the values were partly overlapping we found that contact sites were significantly wider on *mpk3* stigmas compared with Col-0 (Fig. 7b). This demonstrates that MPK3 contributes to the stigma-pollen interaction and structure.

**Fig.7.**
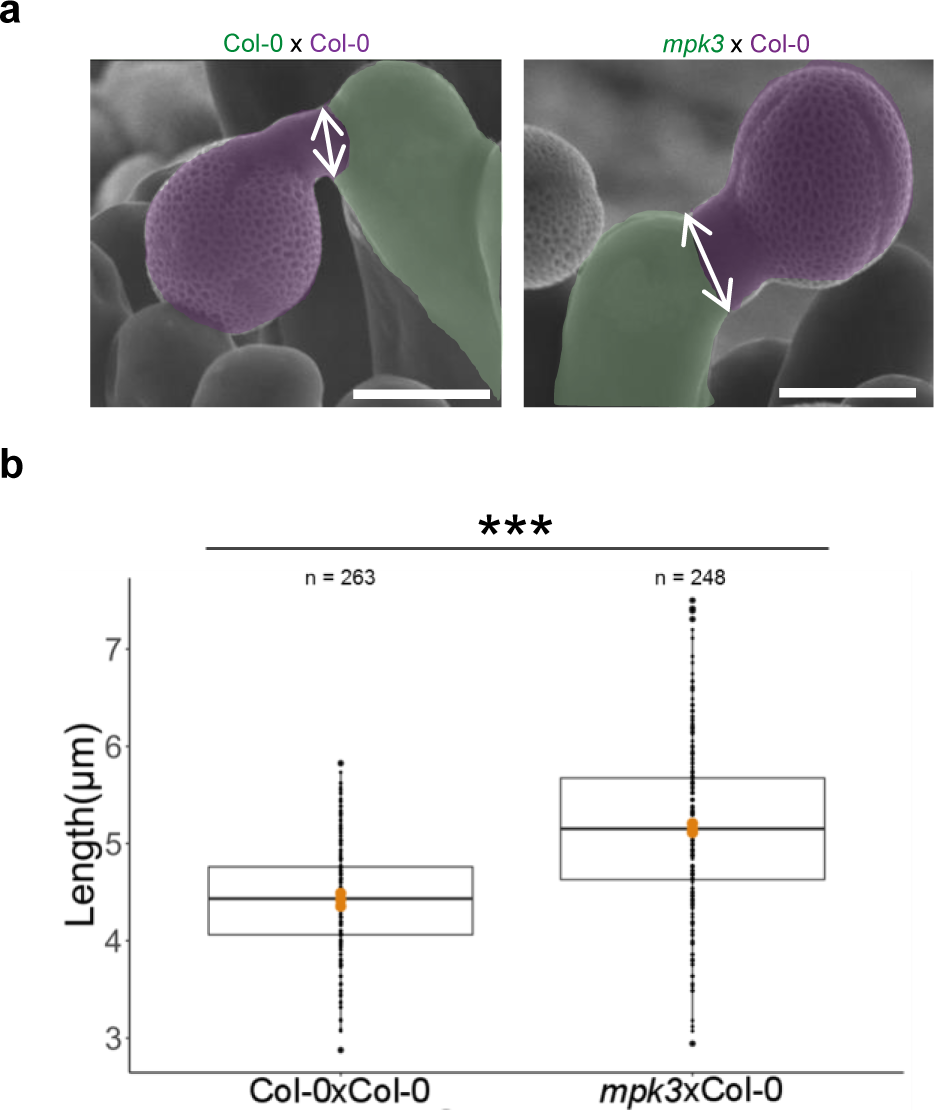
*mpk3* stigma changes the width of pollen tube contact sites. **a**, Pollen tube contact sites on Col-0 (left) or *mpk3* (right) papilla cells observed by SEM 60 min after pollination. One stigma papilla is artificially colorized in green and one pollen grain and its tube in violet, respectively. Scale bar = 10 μm. Double-arrows indicate the width of pollen tube contact site analysed in b. **b**, Width of the pollen tube contact site for each pollination type presented in a. Observations were performed three different days; n is the total number of contact sites measured on 16 Col-0 and 25 *mpk3* stigmas. Orange points in the boxes are averages of each day.*** indicates p-value < 0.01.

## Discussion

To decipher the mechanisms that control reproduction in flowering plants, we extracted the global transcriptomic signatures associated with both pollen and stigmatic cells following their interaction during compatible or incompatible pollination. Although transcriptomes of pollinated stigmas have been recently published^17,18^, these analyses did not try to distinguish between pollen and stigma transcripts. In the present work, we provide a comprehensive transcriptomic dataset for stigma-pollen interactions using a novel SNP-based analysis. We took advantage of a recently developed statistical tool called ASE-TIGAR, which is based on a Bayesian approach to estimate allele-specific expression in diploid cells^21^. Nariai et al. showed an accurate estimation of gene expression from RNA-seq with ASE-TIGAR and identified some autosomal genes as allele-specific genes in a human reference lymphoblastoid cell line. We applied this new tool to discriminate transcripts from female-male mixed tissues during the first steps of the fertilization process. Our experimental design and bioinformatic pipeline allowed us to unveil the transcriptomic response of the pollen/pollen tube and that of the stigma following compatible or incompatible pollination at two early (10 min and 60 min) time points of the male-female interaction (Fig. 1). It is worth noting that another SNP-based pipeline has recently been published^20^, identifying transcripts derived from the pistil and pollen tubes collected 8h after compatible pollination. Although we cannot compare the two analyses directly, as pollination time points were different and only compatibility was studied by Leydon et al., both analyses succeeded in distinguishing male and female transcripts as presented in Fig. 2c. However, while Leydon et al., 2017 used 12 % of reads that had SNPs among the total sequenced reads, in our analysis we took all the RNA-seq reads into account, including those without SNPs^20,21^. This difference in the approaches has likely contributed to making our analysis more exhaustive, as exemplified by the remarkable segregation we found between male transcripts and those derived from female and vegetative tissues (Fig.2 c). The sex preferentially- or specific-expressed gene lists we drew up contain many of the already identified female or male specific genes and reported GO term enrichment^8,13,14^ (Fig.2). Our analysis also clarified the expression of genes that have been reported to have function during pollination. The exocyst complex, including the EXO70A1 protein, has been shown to play a role in pollen acceptance, probably through its function in the secretion of factors required for pollen hydration and pollen tube growth^13,14^. We found *EXO70A1* to be stably expressed in stigmas over time (nFPKM(stigma) = 27.1 at C0, with no expression change following pollination). *ACA13* which was shown to be involved in calcium fluxes was up-regulated after both compatible (FC=3.2 at C60) and incompatible (FC=2.4 at I60) pollinations; similar up-regulation after compatible pollination was also reported by Iwano et al., 2014.

Together with the correlation analysis (Fig. 2c) and experimental evaluations (Fig. 2b) we can conclude that our female-/ male-transcripts globally represent stigma-/pollen-transcripts. Transcriptomic dynamics shown by PCA and Venn diagrams were not substantial in terms of variance and number of DEGs (Fig. 3ab). These results are in line with those of Matsuda et al.^17^ who reported only a few percent of difference between transcripts before and 60 min after pollination for both compatible and incompatible interactions.

While our data are rather consistent with other transcriptomic analyses of pollinated stigmas^8,17,35^ (Fig. 2, 3), the clear distinction we made between male and female transcripts also reveals surprises. The Venn diagram suggested a rapid transcriptomic response in the stigma for compatible (414 up-regulated genes at C10) but not for incompatible pollination (45 up-regulated genes at I10) (Fig. 3b left). Since almost no specific transcriptional induction was detected at I10 (only 4 up-regulated genes), this suggests that key molecules required for the incompatibility reaction are already present in the stigma (Fig.3b left, I10up). This may be consistent with the observation that the pollen rejection response in *A. thaliana* lines exhibiting a restored incompatibility system is extremely rapid and occurs within minutes^12^. Astonishingly, while no cellular changes were detectable by SEM on stigmas one hour after incompatible pollination (Fig. 1a), we identified more than one hundred up-regulated genes in stigmas and only 6 in pollen at I60 (Fig. 3b). This demonstrates that the stigma undergoes major molecular changes following incompatible pollination, i.e. that incompatibility is actively maintained through a global physiological modification of the stigmatic cells. The stigma up-regulated genes can be assimilated to the stigmatic response to self-incompatible pollen. As the incompatibility reaction is maintained for several days, at least as long as SRK is properly expressed in the stigma, we may propose that among the stigma up-regulated genes there are key factors required for the maintenance of SI. Since components of the signaling pathway downstream of the SRK-SP11/SCR receptor-ligand complex are still not fully unveiled, our data provide a key resource to identify new genes involved in the incompatibility response and its maintenance over time.

Our results revealed a molecular signature of compatible pollination in stigma involving the activation of stress response genes and PTI pathways (Figs. 3, 4, 5). The similarity between the stigma-pollen interaction and plant-pathogen interaction has been discussed and has drawn researchers’ attention for a long time^10,43–45^. Several transcriptomic analyses of pollinated stigmas/pistils reported enrichment in transcripts linked to stress or defense response in female tissues^18,20,34,45,46^. Tung et al. and Leydon et al. suggested that such defense-related genes could have functions in pollination. Moreover, although incompatible response has been considered to share common mechanism with plant-pathogen interaction as it is a response to block the invader^18,45^, in our analysis, GO terms associated with defense responses did not clearly appear after incompatible pollination but rather were mainly associated with the compatible response (Fig. 3c).

Recently, Zhang et al.^18^ performed a time course transcriptome analysis of compatible and incompatible reaction in *Brassica napus*. They speculated that compatible responses had close parallels with plant-pathogen interactions, mainly with effector-triggered susceptibility, while incompatible response rather resembled effector-triggered immunity, and that PTI could be common to both compatible and incompatible responses. Although we found up-regulated defense related genes in the incompatible reaction, no clear defense pathways were identified by KEGG analysis. By contrast, our study clearly indicates that two PTI pathways were induced during compatible response (Fig.4). Indeed, the SNP-based analysis demonstrated the stigma specific expression and induction of genes involved in these pathways (Fig. 5). This result was not completely unexpected since pollen tube growth resembles pathogen attack from several angles, for example invasive growth recognition of an external invader, cell wall digestion, and calcium burst^8,47,48^. We suggest that plant responses to pollen and pathogens share conserved molecular mechanisms at early stages of the interactions but that later downstream deviations could lead to different reactions, one inhibiting pathogen invasion, while the other promotes pollen tube growth in the pistil. Because links between symbiotic and pathogen interactions have been already been uncovered^49,50^, we can also speculate that the response of plant cells to symbiotic organisms could share common mechanisms with the response of the stigma to pollen. How these shared modules evolved together whilst maintaining highly specific responses (i.e. pollination, PTI, symbiosis) is an intriguing question for future studies.

MPK3 is one of the key components of PTI signaling pathways^38^, and we confirmed both the induction of genes associated with MPK3 expression, and MPK3 activation itself, in compatible pollinated stigmas (Supplementary Fig. 4, and Fig. 6). The increased pollen-stigma contact sites we observed in *mpk3* mutant suggest that the structural properties of the cell wall in the mutant could somehow be impaired (Fig. 7). Although the role of mechanical signals and cell wall reorganization has been studied in growing pollen tubes *in vitro*^51–53^, very little is known about components involved in pollen tube penetration and elongation within the stigma. It seems likely that major cell wall modifications are necessary to allow pollen tube growth through softening and loosening of the cell matrix^48^. How the pollen tube and stigma papilla contribute to these changes is still unknown even though both male and female partners are likely to contribute. The identification of a MPK3-dependent stigma-pollen interaction structure provides an interesting phenotypic indicator for such future studies.

MPK3 is involved in sensing cell wall modifications and reorganization during development as well as following the perception of external signals like pathogen attacks and wounding by receptor-like Wall-Associated Kinases (WAKs)^54,55^. WAKs bind pectin fragments and oligogalacturonic acids that are generated by pathogen attacks and wounding. In stigmas, two of the *WAK* family members (*AT2G23450* and *AT1G79670*) were stably expressed (nFPKM(stigma) = 11.7 and 12.2, respectively at C0 with no expression change following pollination), and another member, *AT1G21210*, was up-regulated after compatible pollination (FC = 2.3 at C60).

Interestingly, Glycine-Rich Protein3 (GRP3), a possible ligand for WAKs that is known to bind to WAK1^56^ was highly expressed (nFPKM(stigma) = 303.0 at C0 and no expression change following pollination) in stigmas. In addition, we found that several pectin related enzymes were up-regulated in stigmas after compatible pollination. Taken together, we propose that cell wall reorganization that allows pollen tube growth in the stigma could be dependent on MPK3-based pathways (Fig. 8).

**Fig.8.**
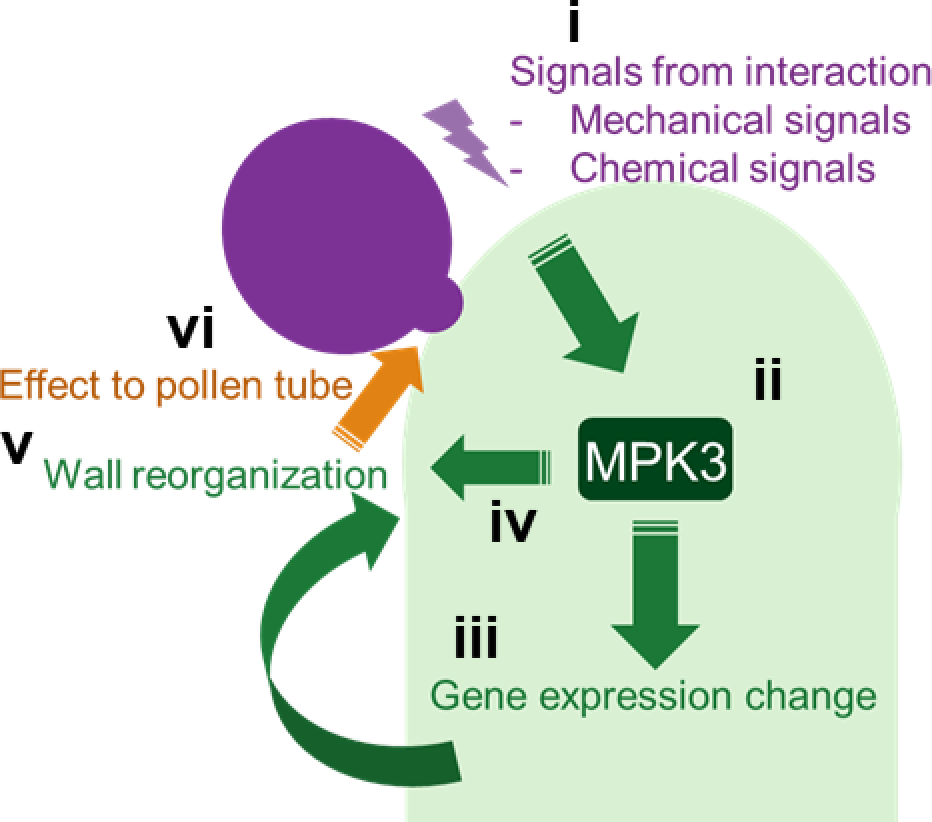
Stigma-pollen interaction involves MPK3-dependent pathway. Schematic model for stigma-pollen interaction. i, Signals at the contact site between pollen tube and papilla cell. ii, Signal transduction involving MPK3 pathways. iii, Gene expression change. iv, Protein activations without transcriptomic change v, Stigma cell wall reorganization. vi, Effect on pollen tube by stigma surface property.

Our study elucidates the molecular signatures of pollen and stigma responses following pollen-stigma interaction in compatible and incompatible situations. Most remarkably, we reveal that the acceptance of pollen grains involves the activation of PTI pathways and we confirm the long-suspected idea that fertilization and plant pathogen responses share common molecular signaling components.

## Materials and Methods

### *A. lyrata SCR14* and *SRK14* gene cloning and plasmid construction

We used Gateway^®^ vectors^57^ (Life Technologies, USA) for expression of transgenes in *Arabidopsis thaliana*. The *AlSRK14* genomic sequence spanning the coding region from the start to the stop codons (3,620 kb) was amplified from genomic DNA of an *Arabidopsis lyrata* individual containing the *S14*-locus with specific AttB-containing primers (5’GGGGACAAGTTTGTACAAAAAAGCAGGCTACCATGAGAGGTGTAATACCAAAGT ACC3’ and 5’GGGGACCACTTTGTACAAGAAAGCTGGGTTTACCGAGGTTCCACTTCCGTGGTGG3’) and subsequently inserted by BP recombination into a pDon207 plasmid. A fragment of 4,081 kb containing the *AlSCR14* gene, 1,844 kb of the 5’ upstream region and 817 bp of the 3’ downstream region were amplified from genomic DNA of *Arabidopsis lyrata* individual containing the *S14*-locus using the AttB-containing primer (5’GGGGACAAGTTTGTACAAAAAAGCAGGCTCGGGTAGCTCAACCTAGCTAAG3’ and 5’ ACCACTTTGTACAAGAAAGCTGGGT CATGATCACCAAAGACAAGATCC3’). This fragment was subsequently inserted by BP recombination into a pDONR-Zeo plasmid. The DNA fragment containing the *Brassica oleracea SLR1* promoter (1.5 kb upstream of the *SLR1* start codon^25,26^) was inserted into a the pDONR P4-P1R plasmid^58^. The *SLR1* promoter, the genomic sequence *AlSRK14* and a 3’mock sequence were inserted in the pK7m34GW destination vectors by three fragment LR recombination. The genomic sequence *AlSCR14*, a 5’mock sequence and a 3’mock sequence were inserted in the pB7m34G. GenBank accession numbers for pg*AlSCR14* (promoter and gene of *AlSCR14*) and p*SLR1*-g*AlSRK14* (*SLR1* promoter and *AlSRK14* gene) are MH680584 and MH680585, respectively.

### Plant material and growth condition

*Arabidopsis thaliana* C24 and Col-0 transgenic plants were generated using *Agrobacterium tumefaciens*-mediated transformation according to Logemann et al. 2006^59^. *AlSRK14* construct was introduced in Col-0 (Col-0/*SRK14*). *AlSCR14* construct was introduced in C24 (C24/*SCR14*). Unique insertion lines homozygotes for the transgene were selected. mpk3-DG^40–42^ was kindly gifted from Marcel Wiermer and Yuelin Zhang. All plants including Col-0 and C24 were grown in growth chambers under long-day cycles (16h light/8h dark at 21°C/19°C).

### Aniline blue staining and observation

Buds at the end of stage 12^60^ were emasculated and 18 hours later, stigmas which have reached the stage 13 or Early 14 (14E), were pollinated with mature pollen grains. After 2 h, pollinated stigmas were fixed in acetic acid 10%, ethanol 50% and stained with Aniline Blue for epifluorescence microscopy observation. Germinated pollen grains with pollen tubes within the stigmas were manually counted. We considered that pollination is incompatible when fewer than 5 pollen tubes were able to overcome the stigmatic barrier^61^.

### Genomic DNA and mRNA preparation

For genomic DNA extraction, about 2 mL volume young inflorescences of Col-0/SRK14 and C24 were harvested, respectively. After grinding the material in liquid nitrogen, DNA was extracted as described previously^62^. For RNA extraction, late stage 12^60^ Col-0/SRK14 flowers were emasculated and 16-20h after emasculation stigmas were pollinated with compatible (C24) or incompatible pollen (C24/*SCR14*). 0, 10 or 60 minutes after compatible pollination (C0, C10 and C60) or incompatible pollination (I10, I60), 50 stigmas were harvested manually using fine tweezers, then frozen in liquid nitrogen and stored at − 80 °C until further processing. After grinding the material in liquid nitrogen, RNA was extracted and purified by using Arcturus^®^ PicoPure^®^ RNA Isolation Kit (Applied Biosystems / Thermo Fisher Scientific) following the manufacturer instruction except that we added a DNAse treatment (Qiagen, catalog#79254). Five biological replicates of RNA at each point were prepared then four replicates were selected for sequencing based on their quality and quantity.

### Whole genome sequencing and variant calling

Library preparation and whole genome sequencing of Col-0/SRK14 and C24 were performed by HELIXIO (Clermont-Ferrand, France; http://www.helixio.com/) with TruSeq^®^ DNA PCR-free Library Preparation kit (Illumina) and NextSeq500 platform (Illumina) applying paired-end sequencing (2×150 bp) (Supplementary Table 1). We then proceeded to the analysis of the sequencing data using GATK3.5 after quality check by FastQC (http://www.bioinformatics.babraham.ac.uk/projects/fastqc), trimming and pairing of the resulting reads using custom Perl scripts. We aligned the reads to the Col-0 reference genome (TAIR10) with BWA^63^, applied GATK^64^ base quality score recalibration, indel realignment, duplicate removal, and performed SNP and INDEL discovery across all samples according to GATK Best Practices recommendations^65,66^. After checking the depth of coverage of two samples by using GATK DepthOfCoverage (Supplementary Fig. 6), we performed filtering applying depth 3 for Col-0/SRK14 and depth 6 for C24 and homozygous for both, then obtained the complete set of variants for each genome as VCF files.

### Production of new reference genomes

Col-0/SRK14 and C24 genome sequences were derived from TAIR10 genome, by introducing the called variants in the sequence, by using GATK FastaAlternateReferenceMaker. Gene annotations from TAIR10 genome were transferred onto the two new genome sequences using RATT^67^ in “Strain” mode and seqret from EMBOSS suite^68^. To characterize polymorphism (SNPs and short indels) between obtained Col-0/SRK14 and C24 genome sequences, inside and outside predicted genes, we aligned chromosome sequences using LAST (v. 938)^69^, after training LAST on chromosomes 1.

### RNA sequencing and expression analysis

Library preparation and RNA sequencing were performed by HELIXIO with TruSeq^®^ Standard mRNA sample Preparation kit (Illumina) and NextSeq500 platform (Illumina) applying paired-end sequencing (2×75 bp). We then proceeded to the analysis of the sequencing data using ASE-TIGAR after quality check by FastQC, trimming and pairing of the resulting reads using custom Perl scripts. First, we derived mRNA reference sequences from new reference genomes and merged them in one FASTA file, then mapped the clean reads on the mRNA reference with Bowtie2^70^. We then run ASE-TIGAR with SAM files produced by mapping and the mRNA reference^21^ (http://nagasakilab.csml.org/ase-tigar/). Raw sequencing data is available at NCBI database (https://www.ncbi.nlm.nih.gov/sra/SRP154565) under SRA accession: SRP154565.

### Comparison with published datasets

To check the consistency of our RNA-seq data with previously published microarray data^32–34^, for each pair of condition, we computed a Pearson correlation coefficient (r) over all genes with SNPs from our variant analysis, using mean of estimated read counts from three or four biological replicates for RNA-seq and mean absolute intensity for multiple microarray replicates. We also computed Pearson correlation coefficients (r) between previously published RNA-seq data^20^ and the microarray data sets.

### Differential gene expression analysis and calculation of nFPKM

Differential expression analysis on the whole transcriptome was performed using DESeq2^27^. After performing principal component analysis (PCA) for all biological replicates at each condition, we removed one replicate at I10 from female samples and one at I60 from male samples, as they were isolated from their other replicates. All the analyses, including the presented PCA, were performed with the sample set after the removal.

Normalized FPKM (nFPKM) was calculated by dividing stigma-or pollen-FPKM by female or male transcript proportion at each condition (C0, C10, C60, I10, I60).

### Criteria to select expressed genes, sex–preferentially or specifically expressed genes

To select expressed genes, sex-preferentially or specifically expressed genes at C0 (Fig. 2a, Supplementary Table 5), we used nFPKM. We defined stigma-expressed genes consisting [nFPKM(stigma) >1 ∩ with SNPs] and stigma-preferentially expressed genes consisting [nFPKM(stigma) > 1 ∩ nFPKM(stigma) ≥ 10 × nFPKM(pollen)]. Sex-specifically expressed genes were selected among sex-preferentially expressed genes applying more stringent criteria. Stigma-specifically expressed genes consist [stigma preferentially expressed genes ∩ nFPKM(pollen) ≤ 1 ∩ nFPKM(stigma) > 100 ∩nFPKM(stigma) > 100 x nFPKM(pollen)], then sorted by nFPKM(stigma) from the largest to the smallest. Vice-versa for pollen (preferentially / specifically) expressed genes

### GO term and enrichment analysis and pathway analysis

GO and gene enrichment analyses were performed using Gene Ontology Consortium website (http://www.geneontology.org/) and PANTHER13.1 software^71,72^. We used “GO biological process complete” for the classification. GO terms with False Discovery Rate (FDR) < 0.05 were considered as significantly enriched.

Pathway analysis was performed using and KEGG PATHWAY Database using KEGG Mapper software (http://www.kegg.jp/kegg/tool/map_pathway2.html)^37^.

### RT-PCR and sequencing

For experimental evaluation of the SNP-based analysis with reverse transcription polymerase chain reaction (RT-PCR) and sequencing, we selected genes among the top-20 specifically expressed genes that have SNPs at suitable positions allowing discrimination of parental origin (stigma vs pollen) (Fig. 2b). RNA at C0 was purified as described above. cDNA was generated using SuperScript^®^ VILO™ cDNA Synthesis Kit (Invitrogen). Primers were designed by Primer3web (http://bioinfo.ut.ee/primer3/) at the flanking region of SNP including sites. PCR was performed with GoTaq (Promega). PCR products were purified using PCR clean-up Gel extraction kit (Macherey-Nagel) and then sequenced.

### Protein extraction and western blotting

Proteins from around 50 pollinated stigmas at C10 and C60 were extracted in SDG buffer (Tris-HCl 62.5 mM pH 6.8, Glycerol 10%, DTT 2%, SDS 2.5 %). Proteins were separated on a 10 % SDS-PAGE gel and immunodetected with Anti-AtMPK3 (Sigma, M8318), anti-phospho p44/p42 MAPK (Cell Signaling, #9101S), or anti-alpha-Tubulin (Sigma, T5168). Band intensities were normalized to tubulin signal using ImageJ software.

### Environmental Scanning Electron Microscopy (SEM)

Late stage 12 stigmas were pollinated with mature Col-0 pollen and 60 min after pollination, stigmas were observed under Hirox SEM (SH-3000 table-top) at −20 °C with an accelerating voltage of 10 kV. The width of each pollen tube–stigma contact site was measured using ImageJ sofware. We applied a Student test to the measurements.

## Acknowledgments

We thank Yvon Jaillais, Olivier Hamant, Gwyneth Ingram for critical reading of the manuscript, SiCE team members for fruitful discussions and comments, Fabrice Besnard for critical reading of the manuscript and for specific advice and script sharing relative to SNP calling, Naoki Nariai for the help to perform analysis with ASE-TIGAR, Alexander R. Leydon for the information for SNP-based RNAseq analysis, Marcel Wiermer and Yuelin Zhang for *mpk*3-DG line, Agnès Attard for the discussion of the bioinformatics data analysis. We gratefully acknowledge support from the PSMN (Pôle Scientifique de Modélisation Numérique) of the ENS de Lyon for the computing resources. This work was supported by Grant ANR-14-CE11-0021.

## Author Contributions

CK, JJ, MDR developed the data analysis pipeline and performed the bioinformatics analyses. AL, JL supported the bioinformatics analyses. AL helped with interpretation of the transcriptomic data. CK is responsible for all experiments and analysis performed in the study except described below. FR produced and characterized Col-0/*SRK14* and C24/*SCR14* lines. CK and LR performed the image acquisition by scanning electron microscope. CK, LR, FR, IFL harvested plant materials and extracted RNA. CK, TG, IFL designed the study and CK, JJ, TG, IFL wrote the manuscript. All the authors contributed to the discussion.

## Additional information

Supplementary Fig. 1

Supplementary Fig. 2

Supplementary Fig. 3

Supplementary Fig. 4

Supplementary Fig. 5

Supplementary Fig. 6

Supplementary Table. 1

Supplementary Table. 2

Supplementary Table. 3

Supplementary Table. 4

Supplementary Table. 5

Supplementary Table. 6

Supplementary Table. 7

Supplementary Table. 8

Supplementary Table. 9

Supplementary Table. 10

Supplementary Table. 11

